# Depletion of resident muscle stem cells inhibits muscle fiber hypertrophy induced by lifelong physical activity

**DOI:** 10.1101/588731

**Authors:** Davis A. Englund, Kevin A. Murach, Cory M. Dungan, Vandré C. Figueiredo, Ivan J. Vechetti, Esther E. Dupont-Versteegden, John J. McCarthy, Charlotte A. Peterson

**Affiliations:** Department of Rehabilitation Sciences, College of Health Sciences, University of Kentucky, Lexington, Kentucky, USA; Center for Muscle Biology, University of Kentucky, Lexington, Kentucky, USA; Department of Physiology, College of Medicine, University of Kentucky, Lexington, Kentucky, USA

**Author notes:** Corresponding Author Charlotte A. Peterson, PhD Room 439 Wethington Building 900 South Limestone Street Lexington, KY 40536.

**Keywords:** Skeletal muscle, aging, hypertrophy, satellite cells

## Abstract

**Background:** A reduction in skeletal muscle stem cell (satellite cell) content with advancing age is thought to directly contribute to the progressive loss of skeletal muscle mass and function with aging (sarcopenia). However, we reported that the depletion of satellite cells throughout adulthood did not affect the onset or degree of sarcopenia observed in sedentary old mice. The current study was designed to determine if lifelong physical activity would alter the requirements for satellite cells during aging.

**Methods:** We administered vehicle or tamoxifen to adult (5 months old) female Pax7-DTA mice for 5 consecutive days to effectively deplete satellite cells. Following a 2-month washout period, mice were assigned to physically active (free access to a running wheel) or sedentary (locked running wheel) conditions. Thirteen months later, at a mean age of 20 months, mice were sacrificed for subsequent analysis.

**Results:** Satellite cell depletion throughout adulthood negatively impacted physical function and limited muscle fiber hypertrophy in response to lifelong physical activity. To further interrogate these findings, we performed transcriptome-wide analyses on the hind limb muscles that experienced hypertrophic growth (plantaris and soleus) in response to lifelong physical activity. Our findings demonstrate that satellite cell function is muscle type-specific; fusion with fibers is apparent in oxidative muscles, while initiation of Gα_i2_ signaling appears to require satellite cells in glycolytic muscles to induce muscle growth..

**Conclusions:** These findings suggest that satellite cells, or their secretory products, are viable therapeutic targets to preserve physical function with aging and promote muscle growth in older adults who regularly engage in physical activity.

## Introduction

It is widely accepted that the reduction in satellite cell content and activity with advancing age directly contribute to sarcopenia (the age-related loss of skeletal muscle mass and function) (1, 2). However, using the Pax7-DTA mouse which enables tamoxifen-inducible satellite cell depletion throughout adulthood, we reported that significantly reducing satellite cell content at 4 months of age did not influence the onset or progression of sarcopenia in sedentary mice as they aged (3). The current study was designed to determine if lifelong physical activity (voluntary wheel running) would alter the impact of satellite cell depletion on muscle aging.

Considerable hypertrophic growth and metabolic adaptations can occur in adult mouse muscle in the absence of satellite cells (4–6). Many signaling cascades have been identified that promote muscle adaptation, among the most studied are the IGF1/PI3K/AKT hypertrophy signaling pathways and AMPK/PGC-1α metabolic signaling pathways. More recently, G protein coupled receptors (GPCRs) and specific G proteins have emerged as mechanisms regulating muscle adaptation (7–10). We aimed to determine if the molecular mechanisms regulating muscle response to lifelong running would differ with and without satellite cell participation, enabling the identification of satellite cell-dependent processes in muscle response to physical activity. Further, we aimed to determine if lifelong physical activity would rescue the aberrant muscle spindle characteristics observed with the depletion of satellite cells (5).

Our results revealed that while satellite cell content did not influence muscle size in sedentary mice or fiber type distribution in response to physical activity, satellite cell depletion limited muscle fiber hypertrophy induced by lifelong physical activity. Further, Satellite cell depletion reduced muscle spindle size and physical function, independent of physical activity. Taken together, these findings suggest that satellite cells are a viable target to delay or offset age-related declines in physical function and promote muscle growth in physically active older adults.

## Materials and methods

### Animals

The Pax7CreER/+-R26RDTA/+ strain, called Pax7-DTA, was generated by crossing the Pax7CreER/CreER mouse strain generated by the Kardon laboratory with the Rosa26DTA/DTA mouse strain(11). The Pax7-DTA mouse allows for the specific and inducible depletion of satellite cells upon tamoxifen treatment, through *Cre*-mediated recombination of the diphtheria toxin A gene in Pax7-expressing cells, effectively killing satellite cells. All animal procedures were conducted in accordance with institutional guidelines approved by the Institutional Animal Care and Use Committee of the University of Kentucky.

### Experimental design

Thirty-nine adult (5-months old) female Pax7-DTA mice were treated via intraperitoneal injection with vehicle (15 % ethanol in sunflower seed oil) or tamoxifen at a dose of 2.5 mg/day for five days, as previously described (4). Following a 2-month washout period, mice were singly housed and randomly assigned to the sedentary group (cage with locked running wheel) or the running group (free access to running wheel) at 7-months of age (n=9-10 per group). A mechanical counter was used to record wheel rotations and was connected to a desktop computer via ClockLab software (Actimetrics, Wilmette, IL). The animals had access to food and water ad libitum and were checked daily for health and wellness. Mice were sacrificed 13 months later, at a mean age of 20 months. Running wheels were locked 48 hours before sacrifice and mice were fasted overnight. Hind limb muscles (plantaris, soleus, TA and EDL were harvested and prepared for immunohistochemical analysis and the contralateral limb for RNA extraction, and then stored at −80 °C. Five mice (3 vehicle-treated and 2 tamoxifen-treated) died over the duration of the study and upon sacrifice several tumors were found in an additional mouse (vehicle-treated) that was excluded from all subsequent analyses.

### Immunohistochemistry

Immunohistochemical (IHC) analysis was performed as previously described (6). Briefly, hind limb muscles were extracted and weighed, then were pinned to a cork block at resting length and covered with Tissue Tek Optimal Cutting Temperature compound (Sakura Finetek, Torrance, CA, USA), and quickly frozen in liquid nitrogen cooled isopentane and stored at −80°C. Frozen muscles were sectioned at −23°C (7 μm), air-dried for at least one hour, and then stored at −20°C. For determining fiber type distribution, muscle fiber CSA, fiber type-specific CSA, and myonuclear density, cross sections were incubated overnight in a cocktail of isotype specific anti-mouse antibodies for MyHC I (1:75, IgG2B, BA.D5), MyHC IIa (neat, IgG1, SC.71), and MyHC IIb (neat, IgM, BF.F3) from Developmental Studies Hybridoma Bank (DSHB, Iowa City, Iowa, USA), along with an antibody against dystrophin (1:100, ab15277, Abcam, St. Louis, MO, USA). Sections were subsequently incubated with secondary antibodies (1:250, goat anti-mouse IgG2b Alexa Fluor 647, #A21242; 1:500, IgG1 Alexa Fluor 488, #A21121; 1:250, IgM Alexa Fluor 555, #A21426) from Invitrogen (Carlsbad, CA, USA), along with the secondary antibody for dystrophin (1:150, anti-rabbit IgG AMCA, CI-1000, Vector), diluted in PBS. Sections were mounted using VectaShield with DAPI (H-1200, Vector). For an additional measure of myonuclear density, soleus cross sections were stained for PCM1, as previously described(12).

Detection of Pax7+ cells was performed as previously described (4). Briefly, sections were fixed in 4% paraformaldehyde (PFA) followed by antigen retrieval using sodium citrate (10 mM, pH 6.5) at 92°C. Endogenous peroxidase activity was blocked with 3% hydrogen peroxide in phosphate-buffered saline (PBS) followed by an additional blocking step with 1% Tyramide Signal Amplification (TSA) blocking reagent (TSA kit, T20935, Invitrogen) and Mouse-on-Mouse blocking reagent (Vector Laboratories, Burlingame, CA, USA). Pax7 primary antibody (1:100, DSHB) and laminin primary antibody (1:100, Sigma-Aldrich) were diluted in 1% TSA blocking buffer and applied overnight. Samples were then incubated with anti-mouse biotin-conjugated secondary antibody against the Pax7 primary antibody (1:1000, 115-065-205, Jackson ImmunoResearch, West Grove, PA, USA) and anti-rabbit secondary for laminin (1:250, A11034, Alexa Fluor 488, Invitrogen, Carlsbad, CA, USA). Slides were washed in PBS followed by streptavidin-horseradish peroxidase (1:500, S-911, Invitrogen) for 1 hour. AlexaFluor 594 was used to visualize antibody-binding for Pax7 (1:100, TSA kit, Invitrogen). Sections were mounted and nuclei were stained with Vectashield with DAPI (H-1200, Vector). Pax7+ satellite cells were identified as Pax7+/DAPI+.

Detection of N-acetyl-d-glucosamine was evaluated on muscle sections using Texas Red-conjugated wheat germ agglutinin (WGA) (eBiosciences, San Diego, CA). Sections were fixed in 4 % PFA and then incubated with WGA conjugate for 2 hours at room temperature.

### Image quantification

All WGA-stained images were captured at ×10 magnification, and the staining was quantified using the thresholding feature of the AxioVision Rel software. The total area occupied by WGA was quantified and in the case of spindle fibers, it was normalized to spindle fiber circumference and reported as the extracellular matrix (ECM) index. Further, WGA images were used to trace the cross-sectional area of spindle fibers and the intrafusal fibers located within the spindle fibers. The average CSA of the intrafusal fibers within a given spindle fiber was averaged and reported as the mean intrafusal fiber area (μm2) per spindle fiber. All remaining images were captured at 20x magnification at room temperature using a Zeiss upright fluorescent microscope (Zeiss AxioImager M1 Oberkochen, Germany). Whole muscle sections were obtained using the mosaic function in Zeiss Zen 2.3 imaging software. To minimize bias and increase reliability, fiber type distribution, muscle fiber CSA, fiber type-specific CSA, myonuclear density and fiber type-specific myonuclear density were quantified on cross sections using MyoVision automated analysis software (13). To determine satellite cell density (Pax7+ cells/fiber), satellite cells (Pax7+/DAPI+) were counted manually on entire muscle cross sections using tools in the Zen software. Satellite cell counts were normalized to fiber number, delineated by laminin boundaries. All manual counting was performed by a blinded, trained technician.

### RNA isolation

Soleus and plantaris muscles were homogenized in QIAzol (Qiagen, Hilden, Germany) and RNA was isolated using RNeasy Mini Kit (Qiagen) according to the manufacturer’s instructions. RNA was eluted in nuclease-free water for subsequent microarray and RT-qPCR analyses. RNA quantity and purity was checked by measuring the optical density (260 and 280 nm) using Nanodrop. All 260/280 ratios were above 1.8.

### RT-qPCR

Complementary DNA was synthesized from 500 ng of total RNA using the SuperScript IV VILO cDNA Synthesis Kit (Invitrogen, Carlsbad, California). RT-qPCR was performed on a QuantStudio 5 Real-Time PCR System (Thermo Fisher Scientific, Carlsbad, California), with Fast SYBR Green master mix (Thermo Fisher Scientific), a 10-fold dilution of cDNA into a 10μl final reaction, using primers against *Gpr4, Gnai2, Dgkh, Prkcsh* and *Vcp* (Table S1). RT-qPCRs were performed using the following thermal cycler conditions: 50°C for 2 minutes, 95°C for 2 minutes, 40 cycles of a two-step reaction, denaturation at 95°C for 15 seconds, annealing at 60°C for 30 seconds. RT-qPCR efficiency was calculated by linear regression from fluorescence increase in the exponential phase in the LinRegPCR software v11.1(14). Expression was normalized to *Vcp* using the delta-delta Ct method(15).

### Microarray analysis

The microarray hybridization and processing were performed at the University of Kentucky Microarray Core Facility, according to the manufacturer’s protocol (Affymetrix, Santa Clara, CA). Affymetrix chips (mouse Clariom S array) were used with 50 ng/μl of total RNA derived from a pooled sample of right plantaris and soleus muscles (n=7/8 group). In an additional analysis, we pooled RNA (50 ng/μl) samples from a subset of mice that performed an equal volume of running (n=3/group). We pooled RNA samples based on the experimental results reported by Kendziorski et al. showing that gene expression from RNA pools are similar to averages of individuals that comprise the pool (16).

### Gene expression analysis

Enriched (≥ 1.5 fold) gene expression data sets were transferred to the Venn diagram tool created by the Van de Peer lab (http://bioinformatics.psb.ugent.be/webtools/Venn/). Data sets were then transferred to ConsensusPathDB for over-representation and enrichment analyses, carried out with default parameters (17).

### Functional testing

Functional measurements were performed by a trained technician (blinded to the group assignments) at the University of Kentucky Rodent Behavior Core a week prior to sacrifice.

#### Rotarod test

Sensorimotor coordination was assessed using a rotarod apparatus consisting of a rotating rod suspended 18 inches above a padded floor (San Diego Instruments, San Diego, CA). This system uses a mouse’s natural fear of falling as a motivational tool to test gross motor function. Mice were placed on the rotating rod at a speed of 4 revolutions per minute (rpm) for 60 seconds for their training sessions (two training sessions separated by at least 10 minutes). After successful completion of the training sessions and adequate rest, the speed of the rod was gradually increased to a maximum of 40 rpms for each of the three testing sessions. The trial was complete when the animal fell, or the time period ended (300 second max). Latency to fall (seconds) was recorded, and the average of the three testing trials was reported for each animal.

#### Balance beam

A beam walking protocol was used to evaluate mice on their ability to traverse beams of decreasing widths. The beam walking protocol evaluates motor balance and coordination by assessing both the time it takes the mice to traverse the beam. Following standard acclimatization and training, three beam widths were used to assess balance (28, 17, and 11 mm) and the mice were given 1 min per trial to complete the crossing of the beam to a bedding filled safe-room on the far end of the beam. The mean of the five trials for the 11mm beam was used for analysis. Difficulty traversing only occurred with the 11 mm beam.

### Data Analysis

Results are presented as mean ± SEM. Data were analyzed with GraphPad Prism software via a two factor ANOVA, a two factor repeated measures ANOVA (running over time), or an unpaired Student’s t-test (cross sectional running data). When interactions were detected, post hoc comparisons were made with Sidak post-tests. Statistical significance was accepted at P < 0.05.

## Results

### Satellite cell replete mice (SC+) had higher satellite cell and myonuclear density in response to lifelong physical activity than satellite cell depleted mice (SC-)

Pax7-DTA mice treated with tamoxifen at 5 months of age demonstrated ≥90% depletion of satellite cells (SC-) in hind limb skeletal muscles (plantaris, soleus, tibialis anterior (TA) and extensor digitorum longus (EDL)) at the time of sacrifice at 20 months of age (Figure 1B and C, Figure supplement 1). Free access to a running wheel for 13 months, beginning at 7 months of age, did not result in higher satellite cell number in SC-mice (Figure 1B and C, Figure supplement 1). Contrarily, in satellite cell replete mice (SC+), satellite cell content in both the plantaris and soleus muscles was higher in response to lifelong physical activity, in agreement with the findings of previous work in mice and humans (Figure 1B and C) (18–20). Satellite cell content was not affected by running in the SC+ TA or EDL muscles (Figure supplement 1).

**Figure 1.**
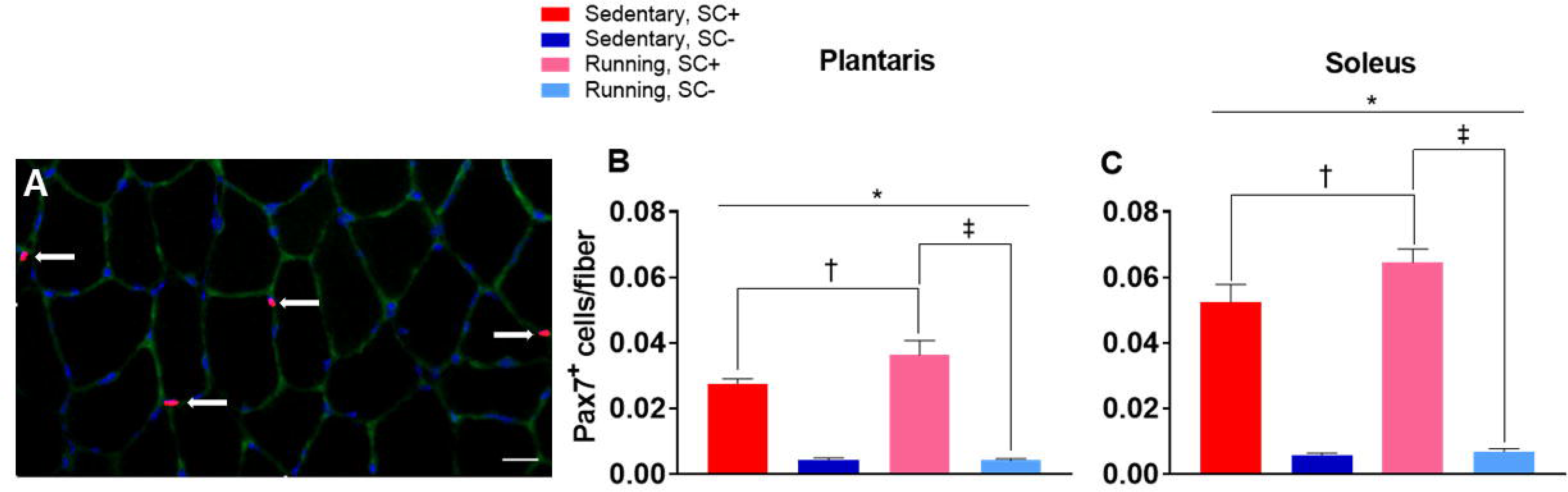
Higher satellite cell density in the plantaris and soleus muscles of running satellite cell replete (SC+) compared to depleted (SC-) mice. (A) A representative image of satellite cell immunohistochemistry showing laminin (green), nuclei (blue), and Pax7 (red; white arrows). Scale bar = 20 μm. (B-C) Satellite cell density in the plantaris and soleus. Data represent mean ± SEM. n = 6-9 mice per group. *P < 0.05, interaction effect between condition and treatment; † P < 0.05, running SC+ vs sedentary SC+; ‡ P < 0.05, running SC+ vs running SC-.

Lifelong physical activity caused higher myonuclear density in the soleus, but not the plantaris, of SC+ mice; as expected there was no change in myonuclear density in these muscles in SC-mice (Figure 2C and D, Figure supplement 2). This finding supports previous work that has indicated the reliance on myonuclear addition during muscle growth is greater in oxidative than glycolytic muscles (19, 21–23). Further, the observation that myonuclear density remained unchanged in the soleus of SC-runners suggests that an alternative stem cell population did not substitute for satellite cells in contributing myonuclei in response to lifelong physical activity. To confirm the higher myonuclear density in the soleus of SC+ mice following running, we performed PCM1 staining to definitively identify myonuclei (12). While, as expected, the PCM1 staining identified more myonuclei compared to DAPI/dystrophin staining, the change in myonuclear density was the same between the two methods (Figure 2C) (12). Similar to satellite cell content, there was no myonuclear addition in the EDL and TA hind limb muscles in response to running (Figure supplement 2).

**Figure 2.**
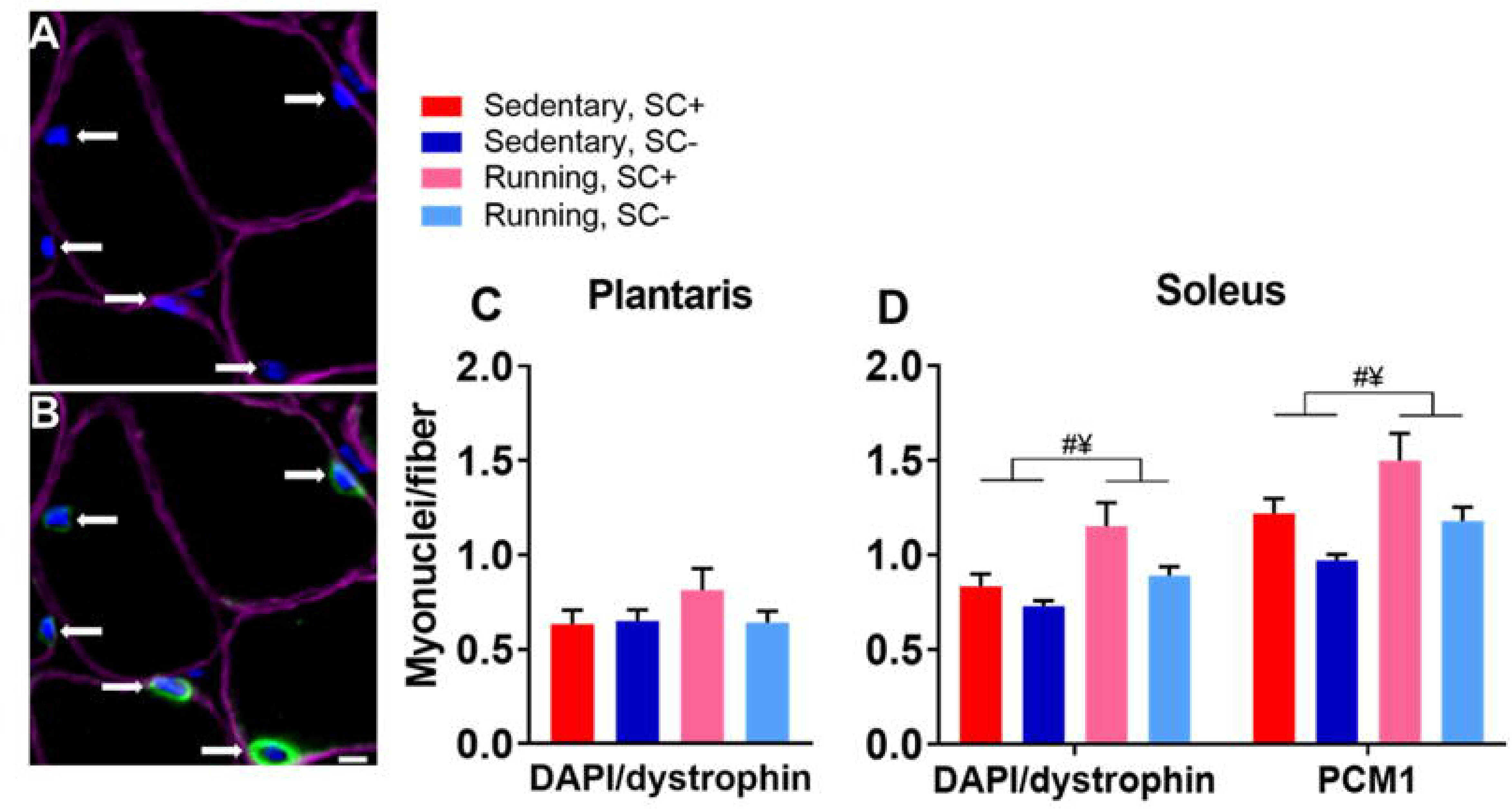
Myonuclear accretion with lifelong physical activity was dependent on the presence of satellite cells. (A) Representative image of dystrophin (purple) and DAPI (blue) staining from muscle cross sections to identify myonuclei (white arrows). (B) Representative image of dystrophin (purple) and DAPI (blue) and PCM1 (green) staining from muscle cross sections to identify myonuclei (white arrows). Scale bar = 5 μm. (C-D) Myonuclear density of the plantaris and soleus. Data represent mean ± SEM. n = 6-9 mice per group. #P < 0.05, running vs sedentary mice; ¥ P < 0.05, SC+ vs SC-mice.

### Lifelong physical activity influenced whole-body and heart weight independent of satellite cell content

Lifelong physical activity led to lower body weight (Figure 3A) and higher normalized heart weight (Figure 3B). These findings highlight that lifelong voluntary wheel running was a powerful stimulus capable of inducing positive phenotypic adaptations independent of satellite cell content.

**Figure 3.**
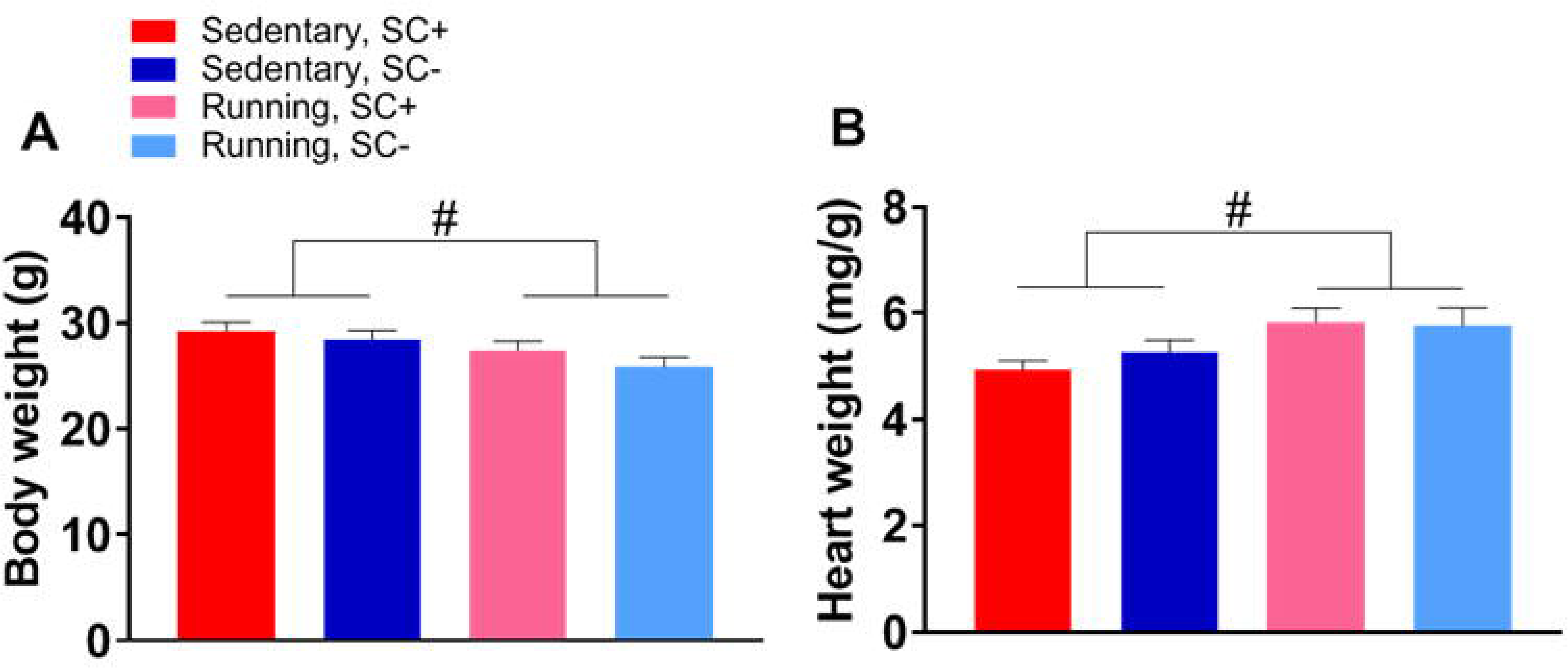
Lifelong physical activity led to a lower body weight and higher heart weight independent of satellite cells. (A) Body weight. (B) Heart weight normalized to body weight. Data represent mean ± SEM. n = 7-9 mice per group. #P < 0.05, running vs sedentary mice.

### Satellite cells were not required for an oxidative shift in fiber type composition in response to lifelong physical activity

In response to lifelong physical activity, immunohistochemical analyses showed a robust shift from glycolytic 2b fibers to oxidative 2a fibers in the plantaris (representative images shown in Figure 4A-B, quantified in Figure 4C). A shift towards an oxidative phenotype in response to increased physical activity is a well reported adaptation in both mice and humans (18, 19, 24). Supporting our previous work following two months of wheel running, we report here that a glycolytic to oxidative shift in the plantaris occurred independent of satellite cell content (Figure 4C) (5). There was no significant shift in the fiber-type composition of the soleus with physical activity, likely due to its inherently oxidative phenotype (Figure 4D).

**Figure 4.**
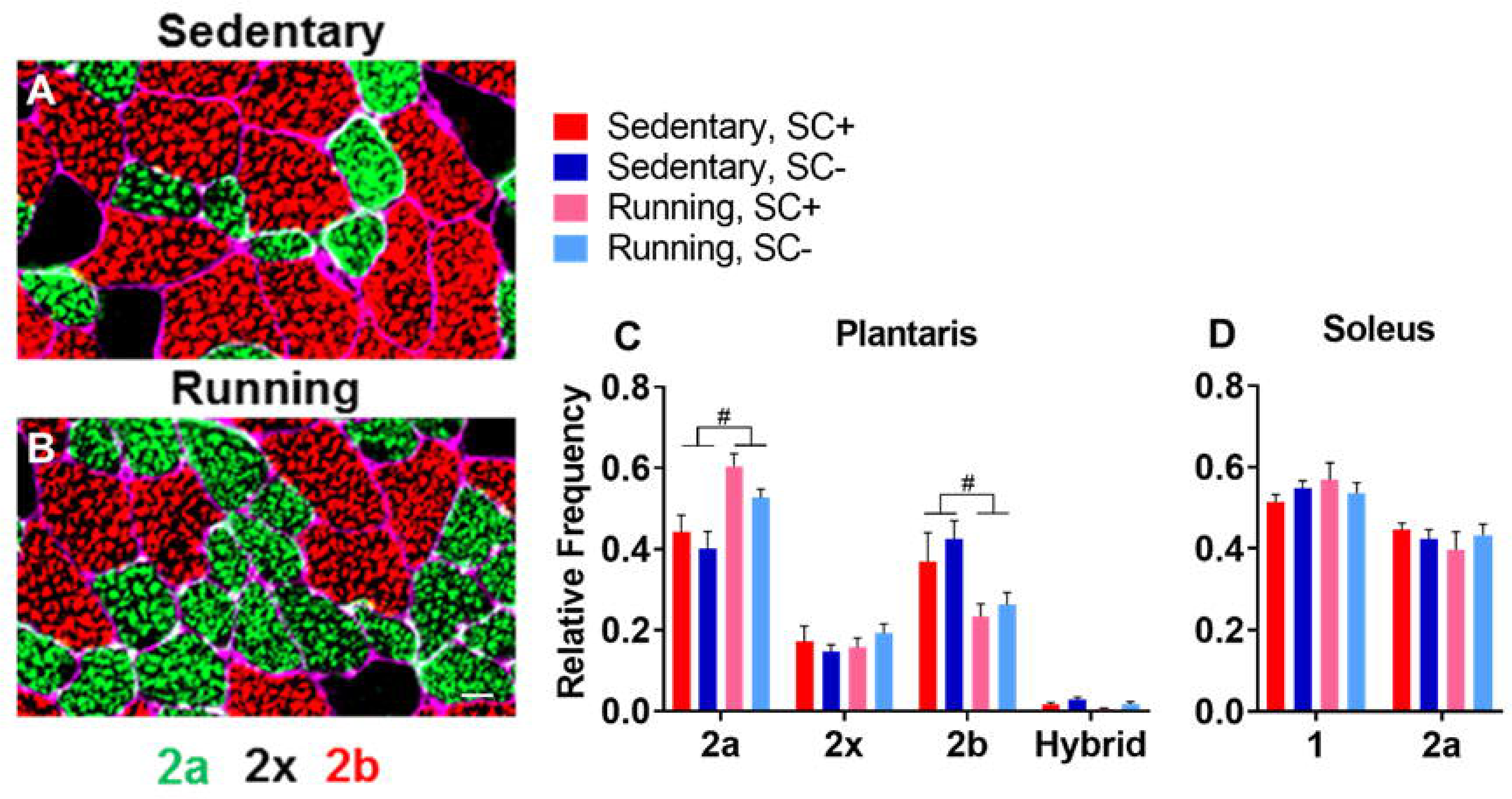
Lifelong physical activity led to an oxidative shift in muscle fiber composition in the plantaris. (A-B) Representative images of the plantaris muscle stained for dystrophin (purple), myosin heavy chain 2a (green) and myosin heavy chain 2b (red). Scale bar = 20 μm. (C-D) Relative frequency of fiber type in the plantaris and soleus. Data represent mean ± SEM. n = 6-9 mice per group. #P < 0.05, running vs sedentary mice.

### Satellite cell depletion limited hypertrophic growth in response to lifelong physical activity

Consistent with our previous report (3), satellite cell depletion in adulthood did not affect plantaris (Figure 5E) or soleus (Figure 5G) muscle fiber cross sectional area (CSA) with age in sedentary mice. Muscle fiber type-specific CSA was larger in plantaris and soleus muscles following lifelong physical activity in SC+ compared to SC-mice (Figure 5E-H). Total mean muscle fiber CSA was larger in the SC+ compared to SC-soleus (Figure 5G); however, due to the prominent shift towards smaller 2a fibers in the plantaris of SC+ runners (as shown above in Figure 4A) mean fiber CSA was not different between sedentary and running mice (Figure 5E and F). Higher muscle fiber CSA with running was a hypertrophic response, as no age-associated atrophy was apparent in the 20 month old mice compared to 8 month old sedentary mice in the plantaris (as reported in Jackson et al. 2015) and the soleus (data unpublished, Jackson et al. 2015 (Figure supplement 3))(5). Greater muscle fiber CSA in response to voluntary wheel running is a common but not universal finding in mice (5, 18, 25, 26). We did not observe a larger fiber CSA with 2 months of running, suggesting that a protracted period of running is necessary to elicit a significant increase in muscle fiber size (5). Lifelong physical activity or satellite cell depletion did not influence muscle fiber size in the TA or EDL (Figure supplement 4).

**Figure 5.**
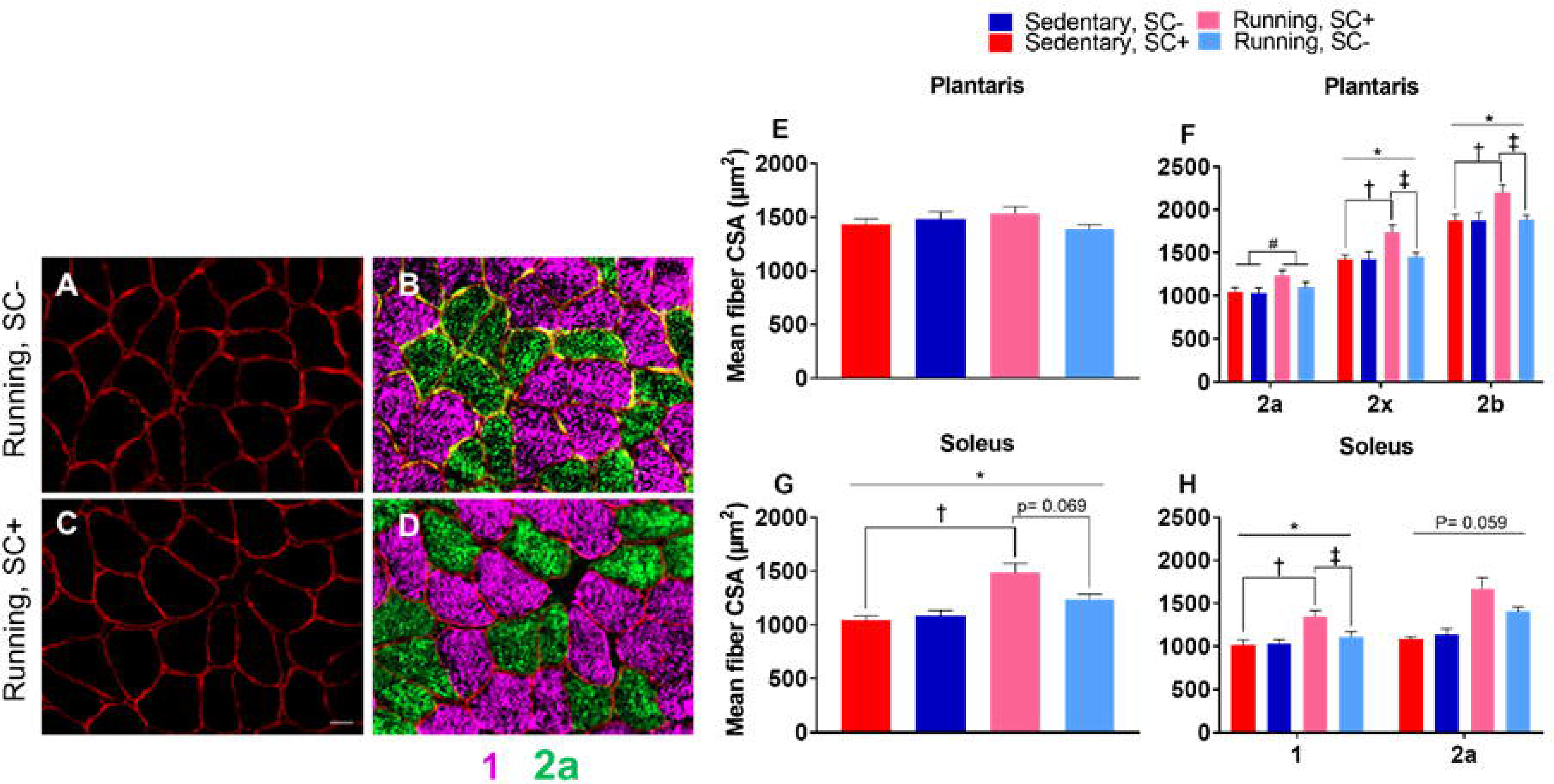
Satellite cell depletion limited the hypertrophic response induced by lifelong physical activity. (A-D) Representative images of muscle fiber size in the soleus stained for dystrophin (red), type 1 myosin heavy chain (pink) and type 2a (green) myosin heavy chain. Scale bar = 20 μm. (E) Whole muscle mean fiber CSA in the plantaris. (F) Mean fiber CSA by fiber type in the plantaris. (G) Whole muscle mean fiber CSA in the soleus. (H) Mean fiber CSA by fiber type in the soleus. Data represent mean ± SEM. n = 6-9 mice per group. *P < 0.05, interaction effect between condition and treatment; † P < 0.05, running SC+ vs sedentary SC+; ‡ P < 0.05, running SC+ vs running SC-.

### Satellite cells influenced the transcriptome-wide response to lifelong physical activity

We next performed transcriptome-wide analysis to identify signaling pathways involved in satellite cell-mediated muscle fiber hypertrophy in response to lifelong physical activity. We pooled RNA samples from the plantaris and soleus (n=7/group) and performed microarray analysis to uncover pathways preferentially enriched in SC+ compared to SC-runners. We compared the enriched genes highly expressed (≥1.5-fold) in the plantaris to those of the soleus and found considerable differences between the muscles in their satellite cell dependent response to lifelong physical activity (Figure 6A). Next, we utilized an over-representation analysis to identify signaling networks significantly enriched in SC+ compared to SC-muscles (Figure 6B and C). We identified a distinct Gα_i2_ signaling pathway with a well-established role in hypertrophic growth to be preferentially enriched in SC+ plantaris (Figure 6D) (9, 10). We validated this finding by performing qPCR on key genes involved in the Gα_i2_ signaling cascade (Figure 6E). Collectively, these data suggest that Gα_i2_ signaling is a potential satellite cell-dependent mechanism stimulating muscle fiber growth specifically in the plantaris in response to lifelong physical activity.

**Figure 6.**
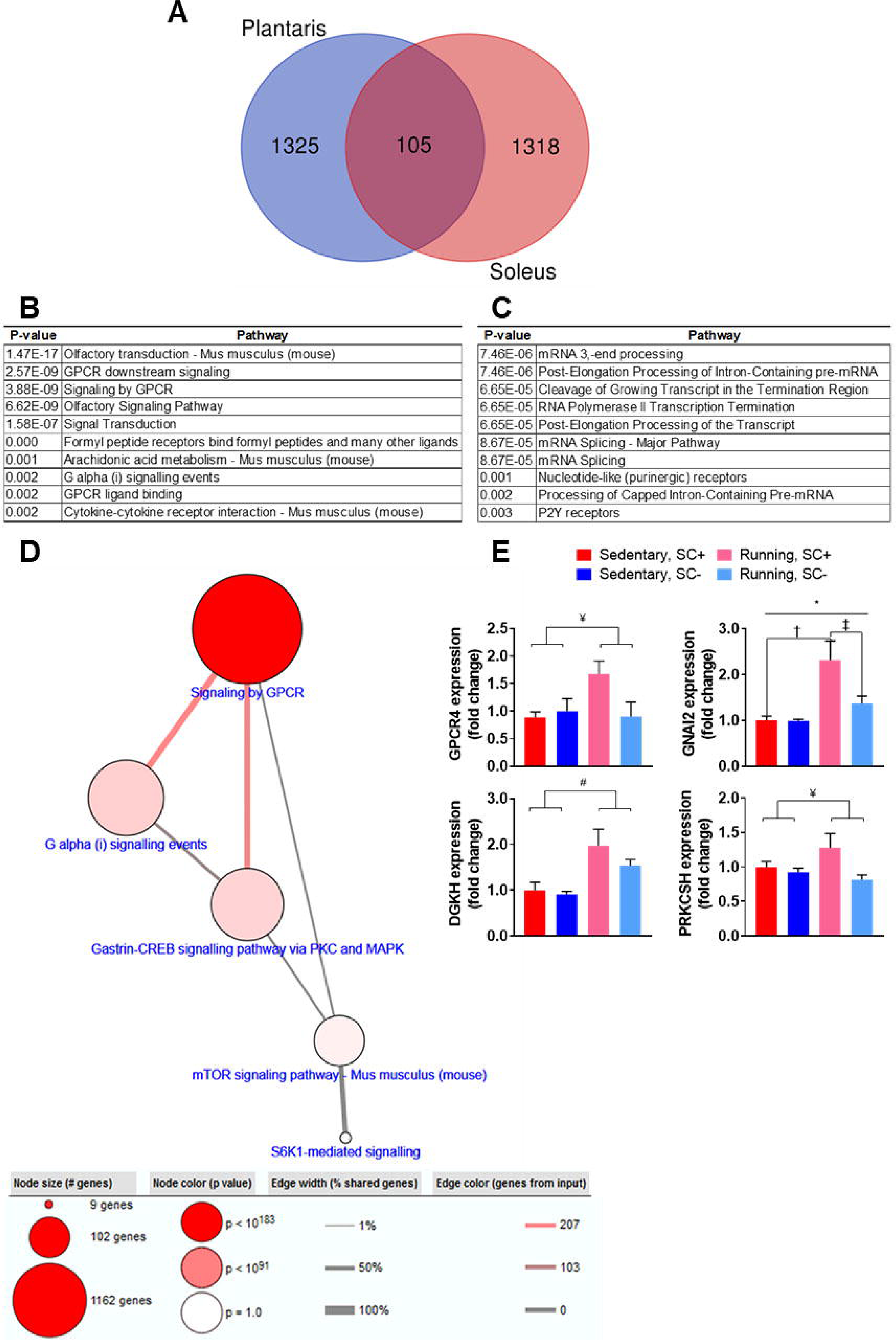
Lifelong exercise in the presence of satellite cells influenced global transcription and up-regulated expression of genes involved in GPCR signaling pathways in the plantaris. (A) Venn diagram of genes enriched in the soleus and plantaris of satellite cell replete runners. Top 10 gene sets overrepresented in the (B) plantaris and (C) soleus. (D) Gene set enrichment analysis highlighting the Gα_i2_ signaling network in the plantaris. (E) RT-qPCR results of Gα_i2_ genes verified the results seen in our gene set enrichment analysis. Data represent mean ± SEM. n = 6-9 mice per group. *P < 0.05, interaction effect between condition and treatment; † P < 0.05, running SC+ vs sedentary SC+; ‡ P < 0.05, running SC+ vs running SC-. #P < 0.05, running vs sedentary mice; ¥ P < 0.05, SC+ vs SC-mice.

### Satellite cell depletion led to aberrant muscle spindle characteristics but not excessive extracellular matrix accumulation

We showed previously that satellite cell depletion throughout adulthood leads to higher extracellular matrix (ECM) accumulation in 24 month old mice, and that ECM accumulation may limit growth in the plantaris during mechanical overload (3, 27, 28). Therefore, we quantified levels of N-acetyl-D-glucosamine utilizing wheat germ agglutinin (WGA) histochemical staining (27). We found no differences in ECM accumulation in the entire cross section (total WGA staining) regardless of satellite cells or running status, ruling this out as a potential mechanism underlying the blunted hypertrophic growth in the absence of satellite cells (Figure 7C).

**Figure 7.**
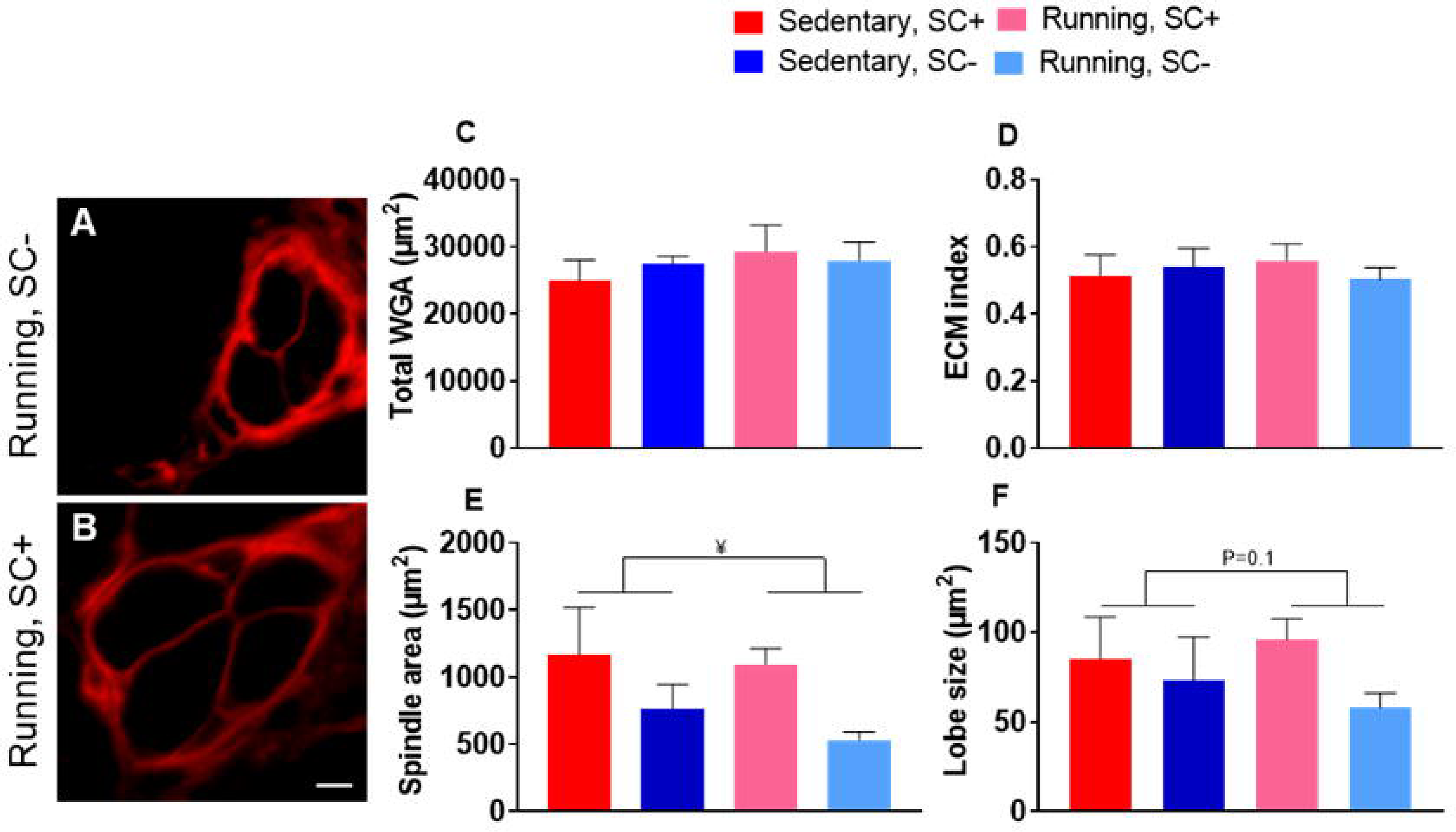
Satellite cell depletion affected muscle spindle characteristics but not ECM accumulation. (A-B) Representative images of muscle spindle fibers from the plantaris muscle. Scale bar = 5 μm. Quantification of (C) total WGA, (D) ECM index, (E) spindle area and (F) spindle lob size. Data represent mean ± SEM. n = 4-8 mice per group. ¥ P < 0.05, SC+ vs SC-mice.

We also reported previously that spindle fiber morphology and function were impaired in satellite cell-depleted muscle, and wanted to determine if lifelong physical activity may rescue this phenotype (5). We found no differences in ECM accumulation within the spindle fibers themselves (ECM index (Figure 7D)). However, satellite cell depleted mice had lower muscle spindle size, as well as lower spindle-lobe size (Figure 7E-F). Lifelong physical activity had no impact on muscle spindle size in SC- or SC+ mice.

### Satellite cell depletion throughout adulthood negatively influenced physical function

As muscle spindles are proprioceptors, we hypothesized that reduced spindle lobe size as a result of satellite cell depletion would negatively impact balance and coordination. To test this hypothesis, we utilized a rotarod test to quantify motor coordination and a balance beam test to assess balance and further assess coordination. SC-mice performed worse on both tests of function, spending less time on the rotarod and taking a longer period of time to transverse the balance beam (Figure 8A and B). These results are similar to those that have been reported previously following 2 months of running in SC-mice and are indicative of poor proprioceptive ability (5, 29). Lifelong physical activity did not improve decrements in performance seen without satellite cells. These findings could, in part, explain declines in physical function with aging that are independent of muscle mass and/or a physically active lifestyle.

**Figure 8.**
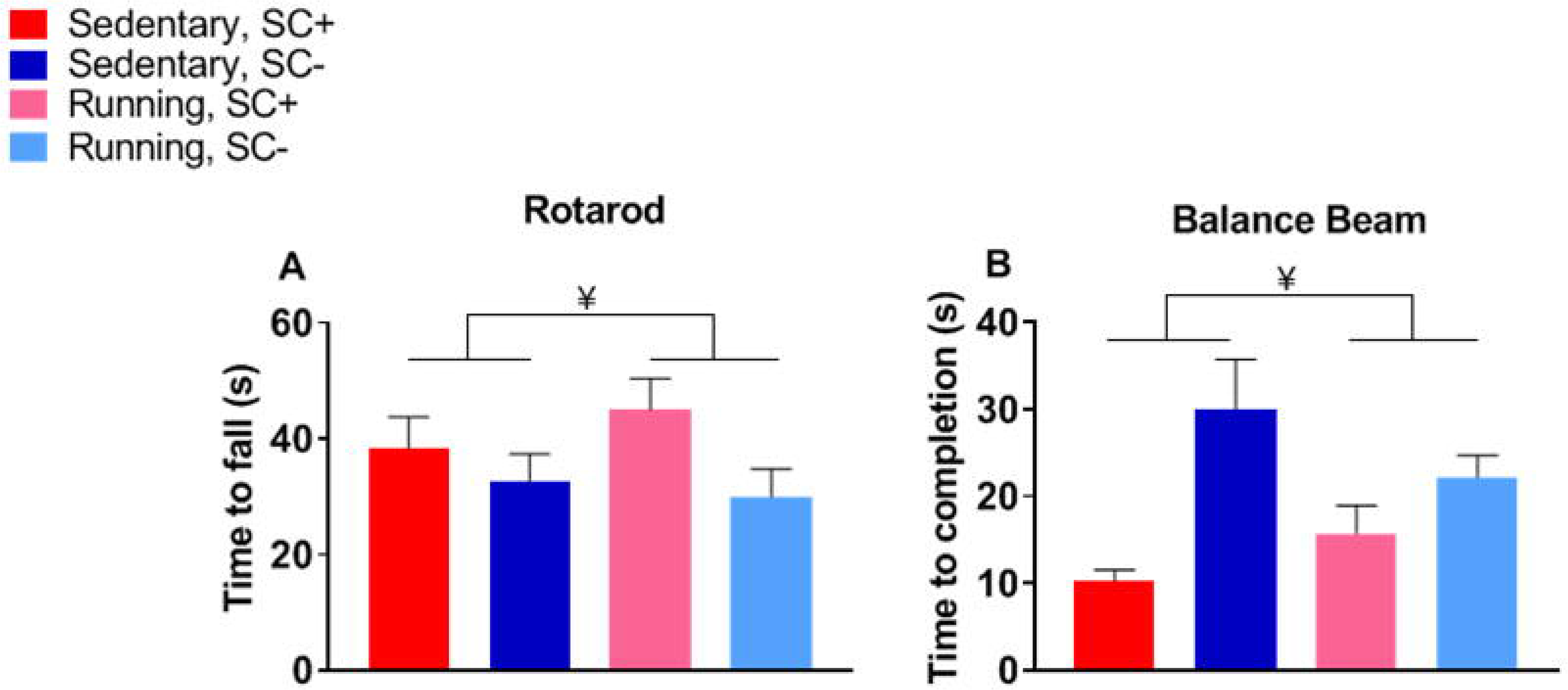
Satellite cell depletion negatively influenced physical function. (A) Time to fall in the rotarod test and (B) time taken to transverse the balance beam. Data represent mean ± SEM. n = 7-9 mice per group. ¥ P < 0.05, SC+ vs SC-mice

### Reduced levels of lifelong physical activity in satellite cell-depleted mice do not explain differences in muscle fiber size

Depletion of satellite cells reduced voluntary wheel running volume (km/day) over the duration of the study (Figure 9A-B). This reduction in running volume was the result of less time spent running (hours/day), as well as slower running speed (km/hour) (Figure 9C-F), likely due to impaired balance and coordination, as shown in Figure 8, above. To determine if differences in running volume affected gene expression and muscle growth, we performed microarray analyses on a subset of SC+ and SC-mice (n = 3/group) that ran equal volumes (∼9 km/day) over the course of the study. We analyzed the genes enriched (≥1.5 fold) in SC+ mice when compared to SC-mice of equal running volume and found that while differences in running volume influenced the transcriptome-wide response of the plantaris (Figure 10A), the most highly enriched pathways and key genes in the Gα_i2_ signaling network appeared to be mediated by the presence of satellite cells, as they were still enriched in SC+ compared to SC-mice when accounting for running volume (Figure 10B-C). Further, differences in muscle fiber CSA persisted in the subset of mice that ran equal volumes over the duration of the study (Figure 10D). These data suggest that satellite cell-dependent Gα_i2_ signaling is critical for muscle fiber hypertrophy in response to lifelong physical activity in the plantaris.

**Figure 9.**
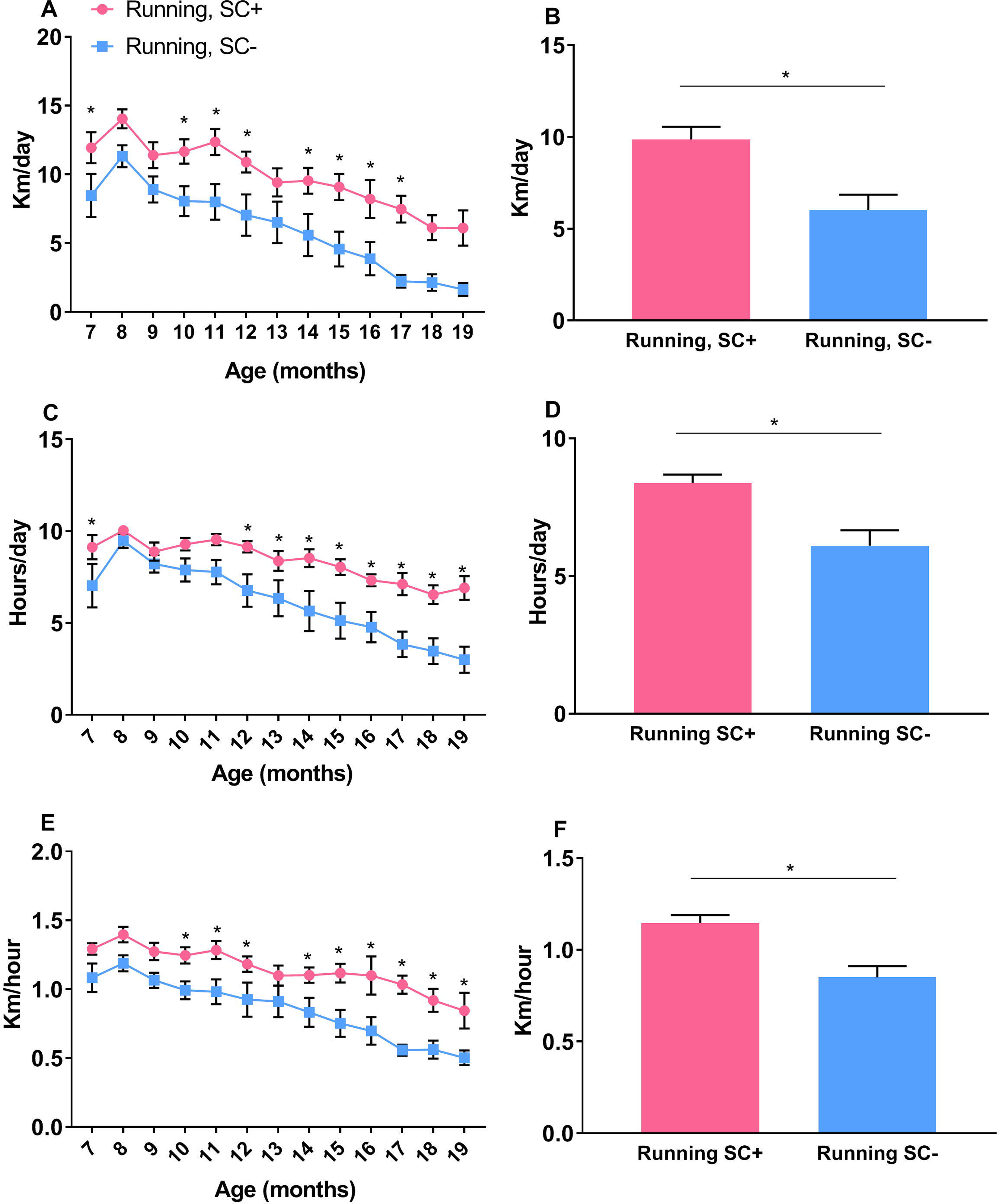
Satellite cell depletion led to reduced voluntary wheel running activity. (A) Average distance run per month over the duration of the study (B) Average distance run over the entire duration of the study. (C) Average time spent running per month over the duration of the study (D) Average time spent running over the entire duration of the study. (E) Average running speed per month over the duration of the study (F) Average running speed over the entire duration of the study. Data represent mean ± SEM. n = 7-9 mice per group.* P < 0.05, SC+ vs SC-mice.

**Figure 10.**
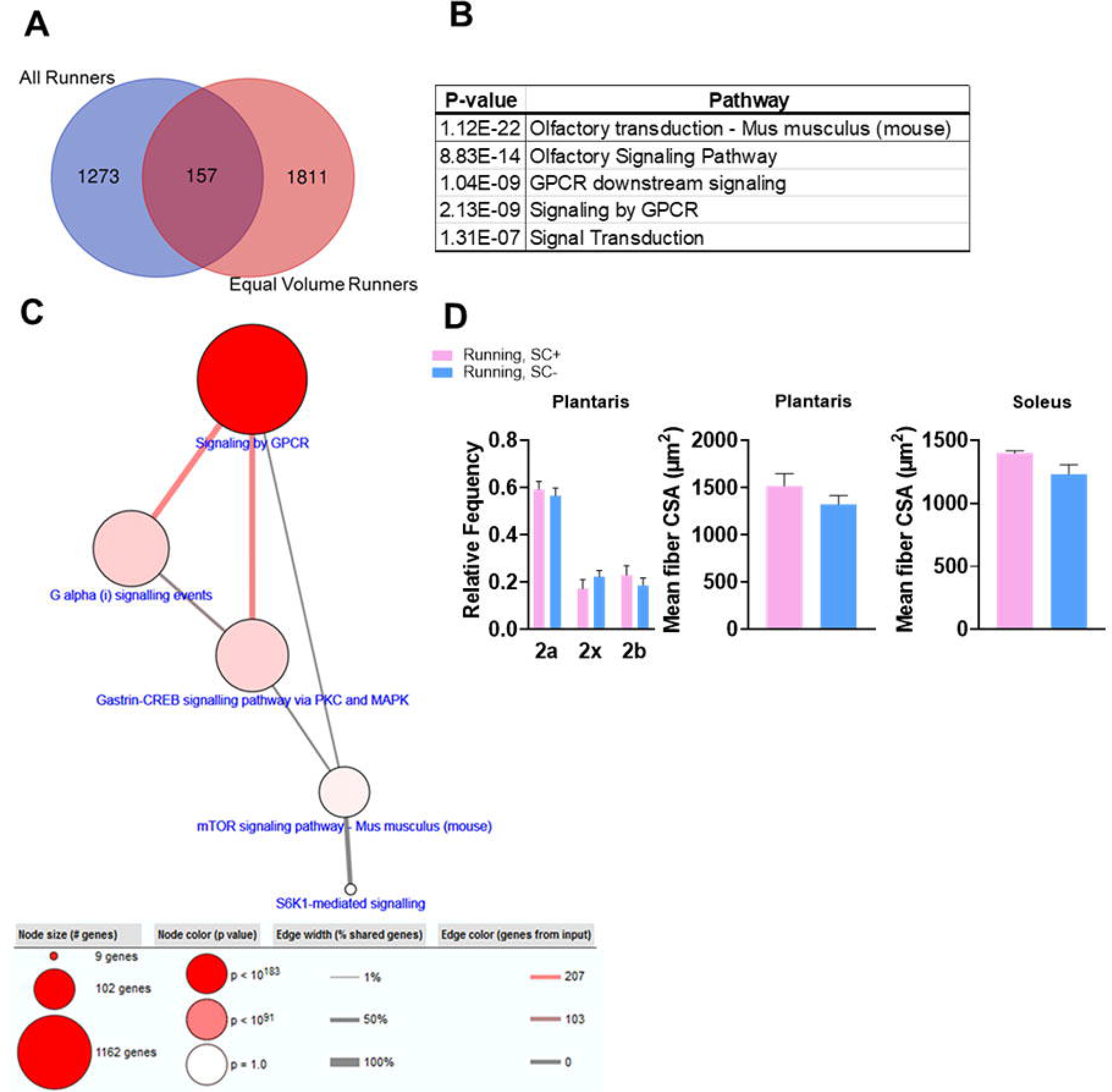
Differences in muscle fiber size and enriched gene sets persisted after controlling for running volume. (A) Venn diagram of genes enriched in the satellite cell replete plantaris of all runners and runners of equal volume. (B) Top 5 gene sets overrepresented in the plantaris of satellite cell replete mice when compared to depleted mice that ran equal volumes. (C) Gene set enrichment analysis highlighting the Gα_i2_ signaling network. (D) Relative frequency of fiber type in the plantaris and mean fiber CSA in the plantaris and soleus. Data represent mean ± SEM. n = 3 mice per group for equal runners; n = 7 for all runners (Venn diagram only).

## Discussion

We report for the first time that a lack of satellite cells throughout adulthood limits muscle fiber hypertrophy in response to lifelong physical activity. Further, satellite cell depletion adversely affects physical function (balance and coordination), as well as running volume. Our findings in satellite cell-depleted and replete control mice support our previous report demonstrating satellite cell depletion does not influence the onset of sarcopenia under sedentary conditions (3). The results reported here also support our previous work showing that the loss of satellite cells negatively impacts voluntary wheel running and muscle spindle characteristics (5).

Satellite cells are not required for the shift in fiber type in response to wheel running, consistent with our previous findings following 2 months of running (5), demonstrating that processes regulating hypertrophic growth and the metabolic properties of the muscle can be uncoupled. It is not surprising that lifelong physical activity leads to higher heart weight and attenuates age-related gains in fat mass that accompany murine aging, and for the first time we report satellite cell ablation does not influence these long term outcomes in response to physical activity. While wheel running induced robust training adaptations in the soleus and plantaris, the TA and EDL were not influenced by the training stimulus. This is likely due to the different functional demands being placed on the muscles via the wheel running stimulus (30) (31).

Larger muscle fiber size in response to lifelong physical activity only in the presence of satellite cells is a clinically meaningful finding. Given the decline in satellite cell content with advancing age, our findings suggest increasing satellite cell content specifically in physically active older adults could be a potential therapeutic strategy to increase muscle mass in response to exercise (32)’(33). Beyond this, falls and fractures are common in older adults, which can result in extended bouts of physical rehabilitation in otherwise sedentary individuals (34). Our results demonstrate targeting satellite cells may augment the hypertrophic response to sustained periods of rehabilitation (5). Further, satellite cell depletion throughout adulthood reduces voluntary physical activity and physical function. Declines in balance and coordination with advancing age increase the risk of falls and are associated with slower walking speeds, a major risk factor for mobility disability and the loss of independence (35–37). We hypothesize that the aberrant proprioceptor phenotype driven by the absence of satellite cells contributes to these outcomes. Whether restoring satellite cell number may rescue the aberrant spindle fiber phenotype and improve physical function remains to be determined, but has important clinical implications.

Satellite cell depletion throughout adulthood does not influence ECM accumulation in sedentary or physically active mice. While the ablation of satellite cells has been shown to induce excessive ECM development in response to mechanical overload as a result of synergist ablation surgery, it is no surprise that the adaptive response to unweighted wheel running is distinct from that of synergist ablation (27). This distinction is likely due to the damaging and supraphysiological nature of synergist ablation-induced muscle overload (38). Based on our previous findings examining the effects of ECM accumulation during sedentary aging in the absence of satellite cells, it appears the magnitude of ECM is highly dependent on the age of the mice being analyzed and the length of satellite cell depletion prior to analysis (3).

We performed transcriptome analysis to identify potential satellite cell-specific mechanisms regulating muscle fiber hypertrophy in response to lifelong physical activity. Our results add to previous work showing distinct mechanisms regulating muscle growth between different muscles and fiber types (21, 39). While satellite cell content is higher in both the soleus and plantaris in response to running, significant myonuclear accretion only occurs in the soleus. It may be that in response to lifelong physical activity, satellite cells potentiate growth in the soleus directly through fusion, whereas they may stimulate growth in the plantaris through signaling to fibers (40). Based on the results of our transcriptome analysis, it appears that satellite cells are critical for the initiation of anabolic signaling events in response to lifelong physical activity and may function via GPCR and downstream Gα_i2_ signaling, specifically in the plantaris. GPCR-mediated signaling events are integral to anabolic signaling and muscle growth (7, 8). Further, the eloquent work of Minetti et al. suggests activation of the Gα_i2_ signaling cascade in the presence of satellite cells likely initiates downstream phosphorylation events necessary for the stimulation of muscle protein synthesis (9, 10). While available deep sequencing data provide insight into the molecular profile of activated satellite cells and information can be gleaned as to the signaling molecules/ligands that are likely candidates influencing GPCR-related signaling, the GPCRs upregulated in the plantaris are orphan receptors, making the identification of bonafide targets challenging (41, 42). Discovering the signaling molecules released by activated satellite cells that lead to meaningful phenotypic adaptations in response to physical activity is a provocative area of study that warrants further exploration.

Our study extends our previous work in sedentary mice by examining the role of satellite cells during aging in physically active mice. Whereas satellite cells do not appear to be required for maintenance of muscle mass in sedentary mice as they age, satellite cells play necessary roles in the maintenance of physical function and in increasing muscle fiber size in response to lifelong physical activity, which may differ by muscle fiber-type composition. In more oxidative muscles, satellite cell fusion may be essential, whereas in glycolytic muscles, satellite cells may promote Gα_i2_ signaling in muscle fibers to facilitate growth. These findings suggest that satellite cells, or their secretory products, could serve as therapeutic targets to preserve physical function with aging and promote muscle growth in older adults engaged in sustained periods of physical activity.

## Acknowledgements

We would like to thank the Rodent Behavior Core at the University of Kentucky and Deann Hopkins for performing functional testing and Dr. Alexander Alimov for overseeing the mice over the study duration. The authors of this manuscript certify that they comply with the ethical guidelines for authorship and publishing in the Journal of Cachexia, Sarcopenia and Muscle.

**Figure supplement 1.** Satellite cells were effectively depleted in the TA and EDL muscles with no effect of running. (A-B) Satellite cell content in the TA and EDL. Data represent mean ± SEM. n = 6-9 mice per group. ¥ P < 0.05, SC+ vs SC-mice.

**Figure supplement 2.** Myonuclear density was not influenced by lifelong physical activity in the TA or EDL. (A-B) Myonuclear density in the TA and EDL. Data represent mean ± SEM. n = 6-9 mice per group.

**Figure supplement 3.** Muscle fiber size was not affected by satellite cell content or lifelong physical activity in the TA or EDL. (A-B) Mean fiber CSA in the TA and EDL. Data represent mean ± SEM. n = 6-9 mice per group.

**Figure supplement 4.** The plantaris and soleus did not demonstrate age-related atrophy. (A-B) Mean fiber CSA in the plantaris and soleus of old (mean age 20 months) sedentary control mice used in this study compared to young (mean age 8 months) sedentary controls; adapted from Jackson et al. 2015.

**Table supplement 1.** qPCR primer list

