## Supplementary figures and images for "Depletion of resident muscle stem cells inhibits muscle fiber hypertrophy induced by lifelong physical activity"

### Fig s1

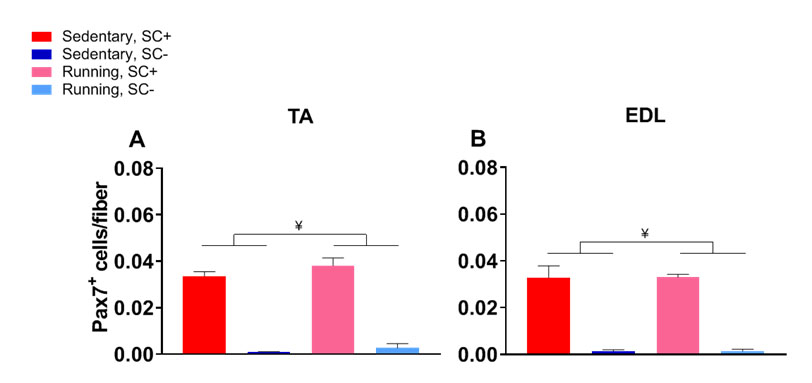

### Fig s2

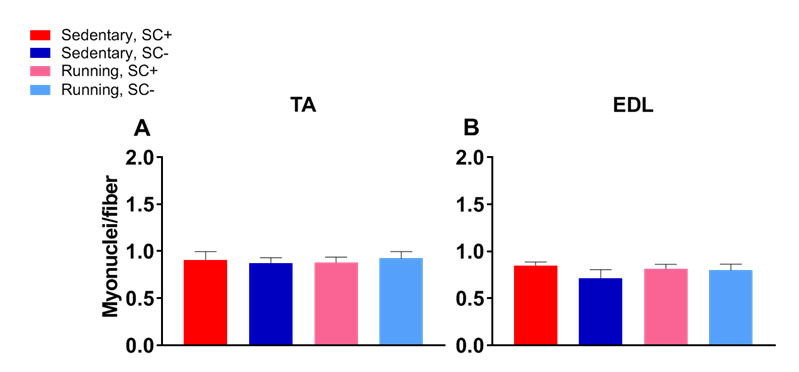

### Fig s3

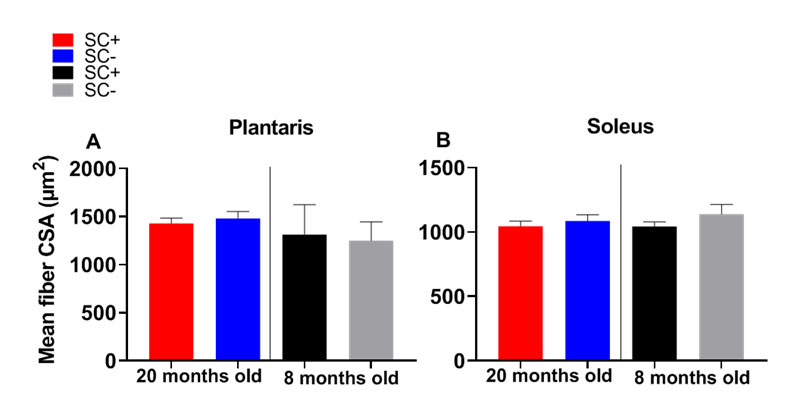

### Fig s4

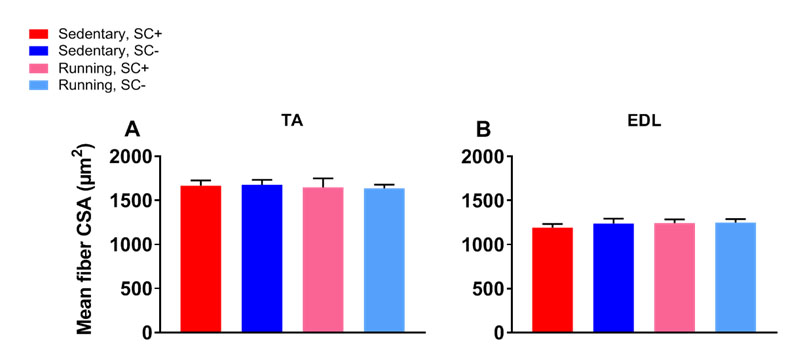
