## Supplementary material for "Depletion of resident muscle stem cells inhibits muscle fiber hypertrophy induced by lifelong physical activity": Table s1

|  |  | **Sequence 5' → 3'** |
| --- | --- | --- |
| Gpr4 | Forward primer | CTGTGCAGAGTCGGGACCAA |
|  | Reverse primer | AGATTGCTAGGCCTGCTCCG |
| Gnai2 | Forward primer | CTGGCAGGATGGGCCAC |
|  | Reverse primer | CATGGTCTTCTTGCCCCCAT |
| Dgkh | Forward primer | CAGCCTGGACCTGGGGATT |
|  | Reverse primer | AAGTTGGGAGGGTTCCGTTC |
| Prkcsh | Forward primer | CGGAGCGTTCCGTTCTCTTA |
|  | Reverse primer | ATGGCCTCTCAGCGAGGTA |
| Vcp | Forward primer | TGCCAGGCCAACTTCATCTC |
|  | Reverse primer | CCGGACATTGGCCTCAGATT |
